# Yes-associated protein (YAP) is required in maintaining normal ovarian follicle development and function

**DOI:** 10.1101/406256

**Authors:** Michele R. Plewes, Xiaoying Hou, Pan Zhang, Jennifer Wood, Andrea Cupp, John S. Davis

## Abstract

Yes-associated protein (YAP) is one of the major components of the Hippo signaling pathway, also known as the Salvador/Warts/Hippo (SWH) pathway. Although the exact extracellular signal that controls the Hippo pathway is currently unknown, increasing evidence supports a critical role of the Hippo pathway in embryonic development, regulation of organ size, and carcinogenesis. The ovary is one of few adult tissues that exhibit cyclical changes. Ovarian follicles, the basic units of ovary, are composed of a single oocyte surrounded by expanding layers of granulosa and theca cells. Granulosa cells (GCs) produce sex steroids and growth factors, which facilitate the development of the follicle and maturation of the oocyte. It has been reported that YAP is highly expressed in human GC tumors, but the role of YAP in normal ovarian follicle development is largely unknown. In current study, we examined YAP expression in bovine ovaries. We demonstrate that downstream hippo signaling effector protein, YAP and transcription co-activator, TAZ, are present and localization of both YAP and TAZ are density-dependent. Likewise, YAP and TAZ are critically involved in granulosa cell proliferation. Furthermore, reducing YAP in granulosa cells inhibits FSH-induced aromatase expression and estradiol biosynthesis. The data suggest that YAP plays an important role in the development of ovarian follicles and estradiol synthesis, which are necessary for maintaining normal ovarian function.

## INTRODUCTION

The ovary is an active endocrine reproductive organ responsible for producing the ovum. During the reproductive lifespan, the ovarian structures and functions undergo continuous change. The most basic unit of the ovary are the follicles. Ovarian follicles play two major functions including steroidogenesis and ovum-oocyte maturation, both required for regulating growth and maintaining environmental homeostasis for reproductive organs. Growth and development of ovarian follicles require a series of coordinated events that induce morphological and functional changes within the follicle, promoting cell differentiation and oocyte maturation. In domestic animals, follicle development and atresia begin during fetal life when primordial follicles are formed. During primordial follicle growth, a single layer of granulosa cells and oocyte develop to primary, secondary and antral follicles. The antral follicle is composed of multiple layers of granulosa cell.

Granulosa cells are principal ovarian cells that undergo dramatic morphological and physiological changes throughout follicular proliferation, differentiation, ovulation, luteinization and atresia. Granulosa cells are responsible for production of sex steroids, enzymes, growth factors, and cytokines required for successful pregnancy. In mammals, follicle stimulating hormone (FSH) released from the anterior pituitary gland, binds to receptors on granulosa cells. Stimulation of FSH leads to the expression of CYP19A1, an enzyme responsible for estradiol biosynthesis [1]. Estradiol released from granulosa cells promotes oogenesis and follicular development. During the estrous cycle, a single dominate follicle develops into a preantral follicle, while many others undergo atresia. Throughout this process, cell proliferation and apoptosis are tightly controlled and synchronized. In human granulosa cells, TGFα, a potent mitogen, has been reported to increase cell proliferation by increasing critical regulators of cell growth and migration [2]. However, despite decades of extensive research, the molecular mechanism of granulosa cells proliferation and apoptosis are still unclear.

The Hippo signaling pathway (also referred to as the Salvador/Warts/Hippo pathway) is highly conserved and plays a critical role in maintain tissue homeostasis by regulating both cell proliferation and apoptosis in various species [3, 4]. This signaling pathway consists of several negative regulators acting in a kinase cascade that ultimately phosphorylate and inactive key downstream effectors and transcription co-activators: Yes-associated protein (YAP) and Tafazzin (TAZ) [5]. The core kinase cascade of the Hippo pathway consists of Ste20-like kinase (Mst1/2) and SAV1 [6]. Mst1/2 are serine/threonine kinases that are enhanced by interaction with the SAV1 scaffold protein. Mst1/2 kinases and SAV1 form a complex to phosphorylate and activate LATS1/2 [7]. Phosphorylated LATS1/2 in turn phosphorylates and inhibits downstream effector protein, YAP and transcription co-activator, TAZ. In the phosphorylated state (inactive), YAP/TAZ are localized in the cytoplasm and sequentially associated with 14-3-3 proteins for degradation [8]. When Hippo signaling is inactivated, this leads to dephosphorylation of YAP/TAZ, allowing for translocation into the nucleus; however, the mechanism is not fully understood. Following nuclear import, YAP/TAZ competes with nuclear VGLL4 for binding with TEAD, a transcription factor, to induce expression of genes that promote cell proliferation and inhibit apoptosis [9].

The Hippo signaling cascade is regulated by numerous upstream signals which include G-protein coupled receptors (GPCRs) [10], Wnt signaling [11], mechanical stress [12, 13], cell polarity [14], as well as microRNAs [15]. Recently, pathways have emerged demonstrating proteins that directly regulate YAP/TAZ activation and localization independent of LATS1/2 kinase activity [16]. Of interest to the ovary, it has long been postulated that extracellular molecules, such as hormones/steroids or growth factors, might regulate the Hippo pathway, controlling cell proliferation and homeostasis [17]. Understanding the influence of FSH and ovarian growth factors, such as TGFα, on Hippo signaling may lead to better understanding of the molecular mechanism of proliferation and apoptosis in granulosa cells.

In reproductive tissues, the exact extracellular signal that controls the Hippo pathway is currently unknown. The ovary is unique in that it is one of few adult tissues that exhibit cyclical changes. Increasing evidence supports that the Hippo pathway plays a critical role in embryonic development, regulation of organ size, and carcinogenesis. Disruption of Hippo pathway has been reported to promote AKT-stimulated ovarian follicle growth [18]. This disruption in the Hippo signaling pathway may result in accumulation of YAP/TAZ in the nucleus, leading to ovarian abnormalities, tumor progression, and impaired fertility. Moreover, it has been reported that YAP is highly expressed in human granulosa cell tumors [19], but the role of YAP in normal ovarian follicle development is largely unknown. The objective of the current study is to determine 1) the localization of YAP and TAZ, 2) the influence of YAP and TAZ in TNF-induced cell proliferation, and 3) the impact of YAP in estradiol production in bovine granulosa cells.

## MATERIALS AND METHODS

### Chemicals

Fetal bovine serum (FBS) was obtained from Valley Biomedical (Winchester, VA). Penicillin-streptomycin and Gentamycin were from Gibco, and Amphotericin B was from MP Biomedical, LLC (Santa Ana,CA). Human FSH was from NHPP/NIDDK (Torrance, CA). DMEM-F12, M199 and other cell culture medium were from Invitrogen (Carlsbad, CA). FBS was from Atlanta biologicals, Inc. (Lawrenceville, GA). YAP, phospho-YAP (ser127) antibodies were from cell signaling technology Inc. (Danvers, MA). Aromatase antibody was purchased from Serotec (Oxford, UK). STAR antibody was obtained from Abcam (Cambridge, MA). topoisomerase II antibody was from EMD Millipore (Billerica, MA), IKBa was from Santa Cruz Biotechnology, Inc (Dallas, TX). Antibodies against β-actin was from Sigma-Aldrich (St. Louis, MO). Second antibodies for Western blotting chemiluminescence were from Jackson Immunoresearch Laboratories Inc. (West Grove, PA); The SuperSignal West Femto Chemiluminescent Substrate Kit was from Pierce/Thermo Scientific (Rockford, IL); Optitran Nitrocellular transfer membrane was from Schleicher&Schuell Bioscience (Dassel, Germany). Nuclear extraction kit was purchased from Active motif (Carlsbad, CA) 3,3-diaminobenzidine (DAB) kit was from Invitrogen (Carlsbad, CA). DAKO LSAB Kit, was from Carpinteria, CA). Mayer’s hematoxylin (MTT (3-(4,5-dimethylthiazol-2-yl)-2,5-diphenyltetrazolium bromide) and antibody against β-actin was from Sigma-Aldrich (St. Louis, MO). YAP1 siRNA was from Dhamarcon/Thermo Scientific (Pittsburgh, PA). [^3^H]thymidine was from MP Biomedicals.

### Bovine granulosa cell isolation

Bovine granulosa cells were isolated from follicles (2-8 mm in diameter) of bovine ovaries. Briefly, GCs were collected by scrapping and flushing the inner layers of follicles. GCs were then washed twice in DMEM-F12 and then plated on the culture dish with 1%FBS or 10%FBS.

### Detection of YAP1 expression in bovine ovarian tissues by Immunohistochemistry

Bovine ovaries were collected during early pregnancy from a local slaughterhouse (JBS^®^ USA, Omaha, NE). Formalin-fixed, paraffin-embedded Ovarian tissue slides were stained with standard streptavidin-biotin immunoperoxidase methods. Briefly, tissue sections were deparaffinized in xylene and rehydrated using a standard alcohol series. Antigen retrieval was conducted by immersing slides in 0.1 M sodium citrate solution and heated to 100°C for 30 mins. Endogenous peroxidase activity was blocked by incubation of slides in hydrogen peroxide (0.3%) in phosphate buffered saline (PBS) for 10 mins at room temperature. The sections were blocked with 10% normal donkey serum at RT for 1 h. The appropriate antibody was applied overnight at 4°C. After washing with PBS, biotinylated secondary antibody and streptavidin peroxidase complex (DAKO LSAB Kit, Carpinteria, CA) were added for 10 mins. After PBS wash, peroxidase activity was visualized with a DAB kit (Invitrogen, Carlsbad, CA). The sections were then counterstained with Mayer’s hematoxylin. Non-immune IgG from the host species was used as a control.

### Subcellular fractionation

Fresh isolated GCs and Cultured GCs (2.5×10^6^) small luteal (2.5×10^6^) were cells collected in PBS with proteinase inhibitors supplied by Nuclear extraction kit (Active motif). Cytoplasmic and nuclear fractions were prepared by following the manufacture’s instruction. 20ug of nuclear or cytoplasmic protein were analyzed by SDS-PAGE and western blot.

### Western blot analysis

Cultured bovine GCs were harvested with ice cold cell lysis buffer (20 mM Tris-HCl (pH = 7), 150 mM NaCl, 1 mM Na2EDTA, 1 mM EGTA, 1% Triton X-100 and protease and phosphatase inhibitor cocktails). The lysed cells were sonicated and cleared by centrifugation at 14,000 × *g* for 5 min. Protein content was determined using a Bio-Rad protein assay kit. The cell lysates (40–60 μg protein per lane) were subjected to 10% SDS-PAGE gel, separated by electrophoresis, and transferred to nitrocellulose membrane as described before (Hou et al. 2010). The membranes were blocked with PBS with 5% BSA and blotted with primary and HRP-conjugated secondary antibodies. Signals were detected using Thermo Scientific SuperSignal West Femto Chemiluminescent Substrate Kit. The imagines were visualized using a Digital Sciences Image Station 440 (Kodak, Rochester, NY).

### Measuring Progesterone level with radioimmunoassay

Conditioned media were collected for progesterone determination using Coat-A-Count Progesterone RIA kit (Siemens, Deerfield, IL) according to the manufacturer’s instructions and as previously reported (Hou et al. 2010).

### YAP knock down

YAP knock down was achieved using siYAP were purchased from Dharmacon. siGLO, (a cy5-labeled non-targeting siRNA as control) from Dharmacon was used as a control. Cells were transfected with either siGLO or siYAP for 6 h using METAFECTENE (Biontex-USA, San Diego, CA) according to the manufacturer’s instruction. After transfection, cells were cultured in DMEM-F12 with 10% FBS for overnight. Cells were then redistributed for cell proliferation assays.

### Cell proliferation assays

Cells were harvested 72 h after siRNA transfection for determination of protein levels or cell numbers. YAP protein expression was determined by western blot analysis and the cell numbers were quantified with an Invitrogen Countess^®^ automated cell counter (Carlsbad, CA).

### [^3^H]Thymidine Incorporation

bGCs (siRNA treated or non-treated) were plated in 24-well plates at a density of 3×10^4^cells/well in DMEM-F12 with 10% FBS. After 24 h, cells were rinsed with PBS and incubated in DMEM-F12 with 2% FBS for 2 h. Cells were then treated with VP with or without TGFa (100ng/ml) for 24 h. [^3^H]thymidine (4 μCi/ml) was added 4 h before the end of the incubation period. Unincorporated radioactivities were removed by washing cells with ice-cold PBS twice followed by the addition of 10% trichloroacetic acid for 10 min at 4°C. Cells were then washed with ice-cold TCA and solubilized with 0.2 M NaOH/0.1%SDS at room temperature. At the end, cells were neutralized with the equal volume of 1N HCL. The radioactivity was determined by liquid scintillation counting.

### Measuring 17β-Estradiol level with Enzyme-linked immunosorbent assay

Conditioned media were collected for 17β-Estradiol determination using an Estradiol EIA kit (Cayman Chemical Company, Ann Arbor, MI) according to the manufacturer’s instructions.

### Microscopy and Analysis

Granulosa cells were seeded at increasing cell densities of 0.125-0.50 × 10^6^ cells/well and maintained for 24 h at 37 °C in an atmosphere of 95% humidified air and 5% CO2. Cells were then fixed with 200 μL of 4% paraformaldehyde and incubated at 4 °C for 30 mins. Cells were rinsed 3 × with 1 mL 1× PBS following fixation. Cells were then incubated with 200 μL of 0.1% Triton-X in 1× PBS at room temperature for 10 min to permeabilize the membranes. Cells were then rinsed 3 × with 1 mL 1× PBS. Cells were then blocked in 1% bovine serum albumin in 1× PBS for 1 h at room temperature. Cells were then rinsed 3 × with 1 mL 1× PBS. Cells were stained with YAP (1:100) or TAZ (1:100) for 1 h at room temperature. Following incubation, cells were rinsed 3 × with PBS to remove unbound antibody. Cells were then incubated with appropriate secondary antibodies at room temperature for 60 min. Cells were rinsed 3 × with 1 mL PBS to remove unbound antibody. Following labeling with antibodies, coverslips containing labeled cells were mounted to glass microscope slides using 10 μL ProLong^®^ Gold Antifade Mountant with DAPI. Coverslips were sealed to glass microscope slides using clear nail polish and stored at −22 °C until imaging.

Images were collected using a Zeiss confocal microscope equipped with a 60× oil immersion objective (1.4 N.A) and acquisition image size of 512 × 512 pixel (33.3 μm × 33.3 μm). The appropriate filters were used to excite each fluorophore and emission of light was collected between 402 to 550 nm. Approximately 30 cells were randomly selected from each slide and 0.33 μm slice z-stacked images were generated from bottom to top of each cell. A 3-dimensional image of each cell was created, and area of individual cell was generated using Zen software. Cells were then converted to maximum intensity projections and processed utilizing ImageJ (National Institutes of Health) analysis software. Corrected total cell fluorescence intensity was determined as previously described [20-22] for the nuclear and cytoplasmic fractions.

### Statistical Analysis

Each experiment was performed at least three times each using cell preparations from separate animals and dates of collection. All data are presented as the means ± SEM. The differences in means were analyzed by one-way ANOVA followed by Tukey’s multiple comparison tests to evaluate multiple responses, or by **t**-tests to evaluate paired responses.

Two-way ANOVA was used to evaluate repeated measures with Bonferroni posttests to compare means. All statistical analysis was performed using GraphPad Prism software from GraphPad Software, Inc.

## Results

### Expression and localization of Yes-associated protein (YAP1) and Tafazzin (TAZ) in bovine granulosa cells (GC) and luteal cells

We first examined YAP and TAZ expression in bovine ovaries using Immunohistochemistry. Our results showed that YAP is largely located in granulosa and theca cells of developing follicles as well as corpus luteum (Figure 1A-C). Very little YAP was detected in stroma cells. YAP is mainly localized in nucleus in granulosa and theca cells (Figure 1A-C), in corpus luteum, however, YAP is mostly occupied cytoplasmic area (Figure 1C). We also checked YAP expression in freshly isolated bovine granulosa cells or plated bovine and large or small luteal cells by western blot analysis (Figure 2). Aromatase and StAR are examined to ensure the purity of the granulosa cell and luteal cell fractions (Figure 2A).

**Figure 1.**
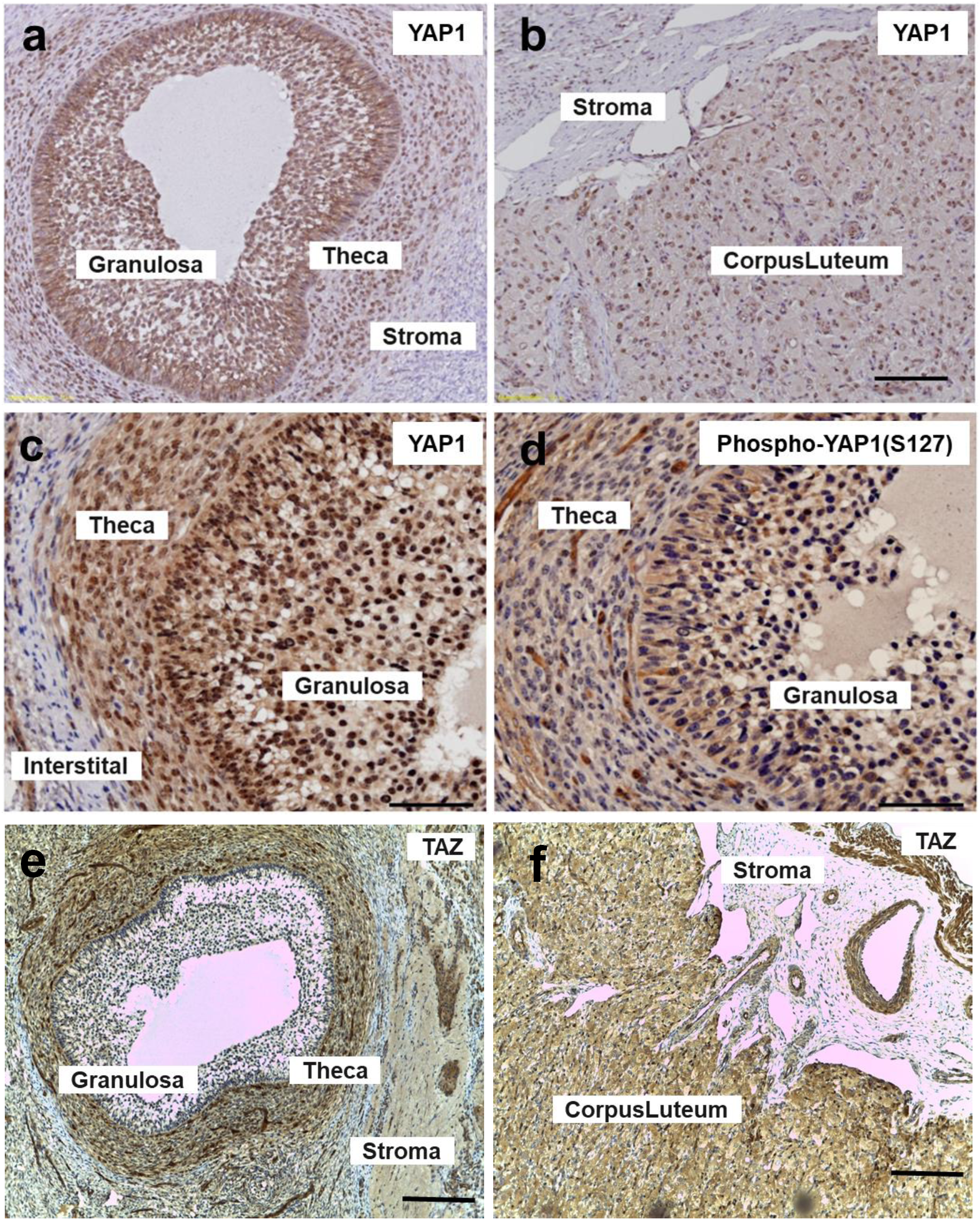
Expression of Yes-associated protein (YAP1) in bovine ovary. Immunohistochemistry was used to determine the presence of phospho-and total YAP1 protein expression in the bovine ovary. Panel A: Representative immunohistochemistry micrograph showing expression of YAP1 in granulosa cells (GC), theca cells (TC). Panel B: Representative micrograph showing expression of YAP1 corpus luteum (CL). Panel C: Representative micrograph showing expression of YAP1 in GC and TC. Panel D: Representative micrograph showing expression of phosphor-YAP1(Ser127) in GC and TC. Panel E: Representative micrograph showing expression of TAZ in GC and TC. Panel F: Representative micrograph showing expression of TAZ in GC and TC. Micron bar represents 500 and 100 μm.

**Figure 2.**
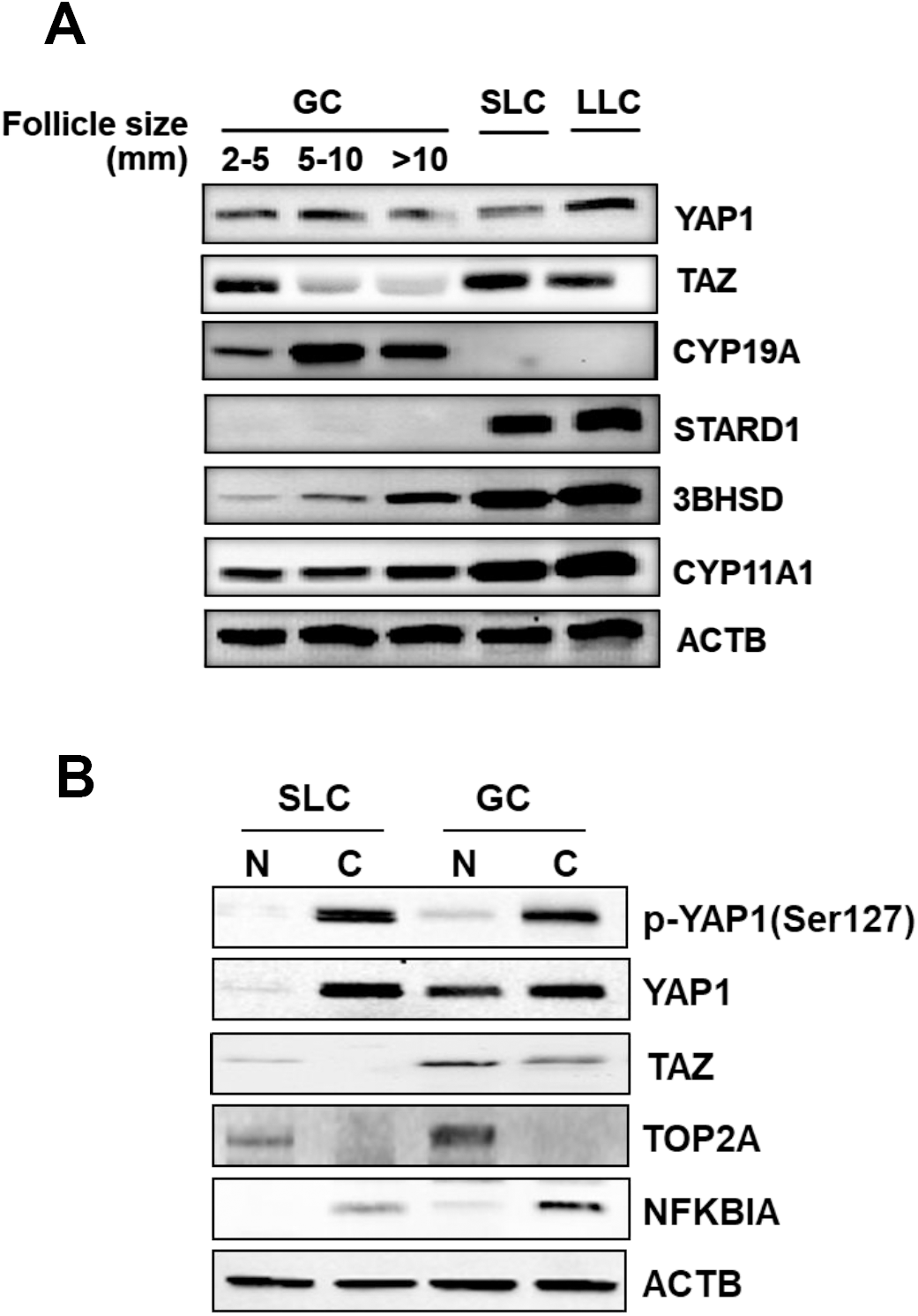
Localization of Yes-associated protein (YAP1) and Tafazzin (TAZ) in bovine granulosa cells (GC) and luteal cells. Panel A: Representative western blot analysis showing Yes-associated protein (YAP1) and Tafazzin (TAZ) expression in freshly isolated granulosa cell (GC), small luteal cells (SLC), and large luteal cells (LLC). Granulosa cells were divided into subpopulations based on follicle size (2-5; 5-10; > 10 mm). Panel B: Representative western blot of YAP1, phosphor-YAP1(Ser127), and TAZ in nuclear (N) and cytoplasmic (C) fractions of GCs and SLCs. Aromatase (CYP19A1; 50 kDa); steroidogenic acute regulatory protein (STARD1; 28 kDa); 3beta-Hydroxysteroid dehydrogenase (3BHSD; 42 kDa); Cholesterol side-chain cleavage enzyme (CYP11A1; 50 kDa); DNA Topoisomerase II Alpha (TOP2A; 190 kDa); Nuclear factor of Kappa light polypeptide gene enhancer in B-Cells 1 (NFKB1A; 39 kDa); Beta-actin (ACTB; loading control; 45 kDa).

### Effects of cell density on localization of Yes-associated protein (YAP1) and Tafazzin (TAZ) in bovine granulosa cells (GC)

Granulosa cells were seeded at increasing cell densities of 0.125-0.50 × 10^6^ cells/well to determine how cell contact/density influenced localization of YAP and TAZ. There was a density-dependent decrease in YAP localized in the nuclear fraction (P> 0.05; Figure 3). Here we show that when cells seeded at the lowest density had 79.6% greater nuclear YAP then when compared to cells plated at 0.50 × 10^6^ cell/well (P> 0.05; Figure 3). Similar to nuclear YAP, cells plated at the lowest density had 67.8% more nuclear TAZ when compared to highest density cells (P> 0.05; Figure 4).

**Figure 3.**
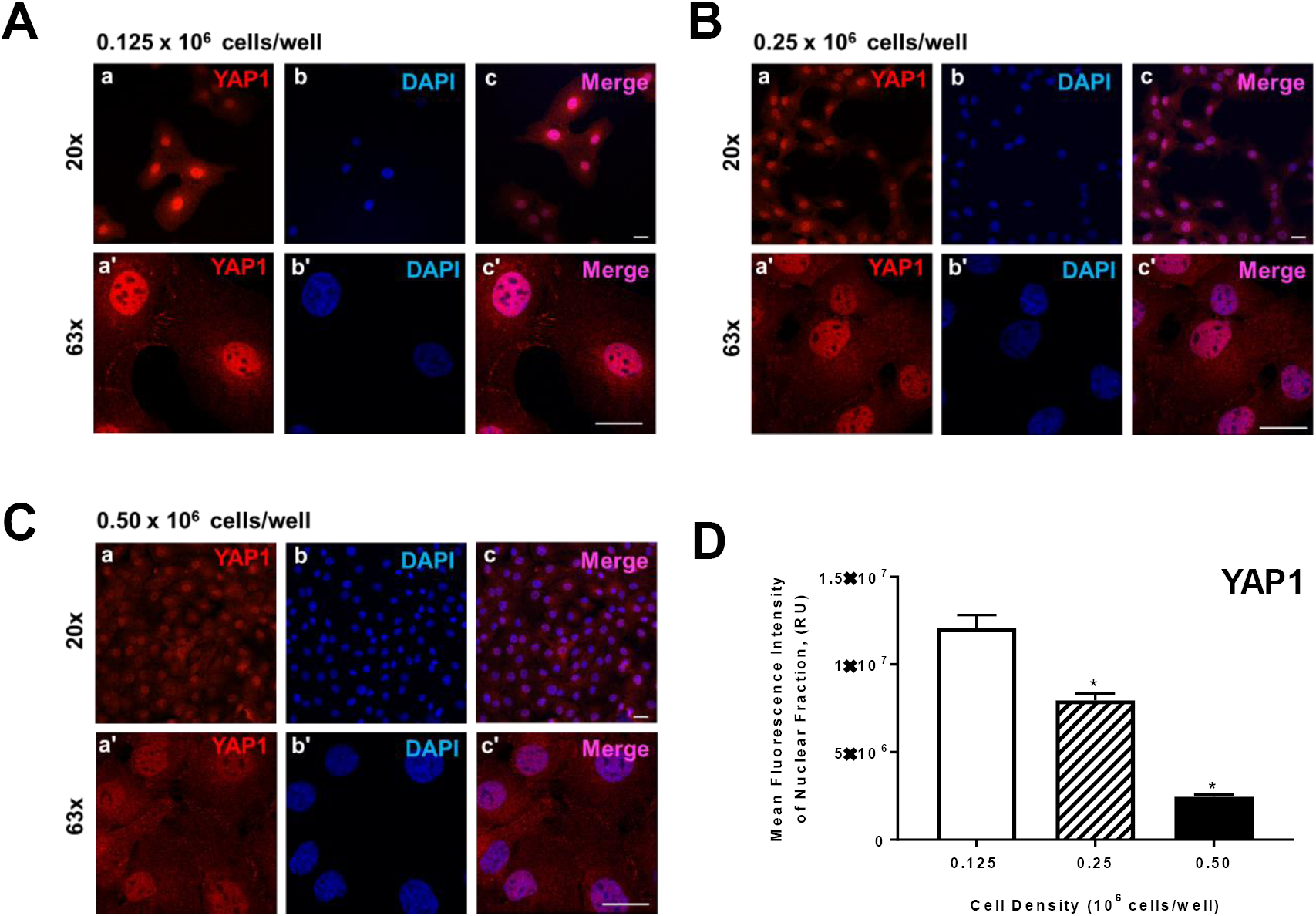
Effects of cell density on localization of Yes-associated protein (YAP1) in bovine granulosa cells (GC). Freshly isolated granulose calls (GC) were seeded in culture dishes at increasing cell densities. Panel A: Representative micrograph of GC plated at 0.125 × 10^6^ cells/well; a) Yes-associated protein (YAP1), 20x magnification, b) DAPI (nuclei), 20x magnification, c) Merge, 20x magnification, a’) YAP1, 63x magnification, b’) DAPI, 63x magnification, c’) Merge, 63x magnification. Panel B: Representative micrograph of GC plated at 0.25 × 10^6^ cells/well. Panel C: Representative micrograph of GC plated at 0.50 × 10^6^ cells/well. Panel D: Quantitative analysis of mean fluorescence intensity of YAP1 expression in nuclear fraction (n = 3). Data are represented as means ± standard error. ^*^Differ significantly within cell density, as compared to 0.125 × 10^6^ cells/well, *P* < 0.05. Micron bar represents 20μm.

**Figure 4.**
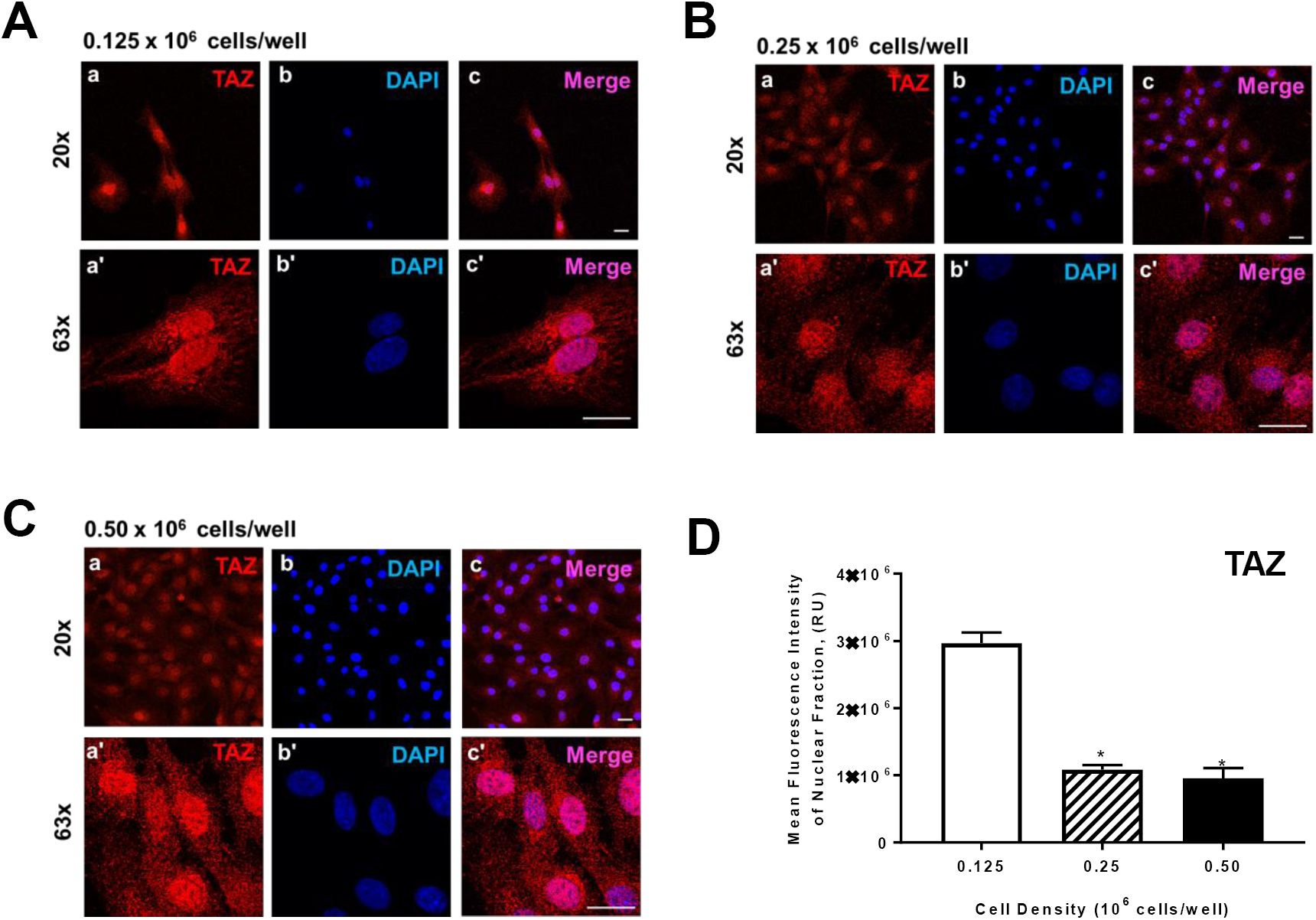
Effects of cell density on localization of Tafazzin (TAZ) expression in bovine granulosa cells (GC). Freshly isolated granulose calls (GC) were seeded in culture dishes at increasing cell densities. Panel A: Representative micrograph of GC plated at 0.125 × 10^6^ cells/well; a) Tafazzin (TAZ), 20x magnification, b) DAPI (nuclei), 20x magnification, c) Merge, 20x magnification, a’) TAZ, 63x magnification, b’) DAPI, 63x magnification, c’) Merge, 63x magnification. Panel B: Representative micrograph of GC plated at 0.25 × 10^6^ cells/well. Panel C: Representative micrograph of GC plated at 0.50 × 10^6^ cells/well. Panel D: Quantitative analysis of mean fluorescence intensity of TAZ expression in nuclear fraction (n = 3). Data are represented as means ± standard error. ^*^Differ significantly within cell density, as compared to 0.125 × 10^6^ cells/well, *P* < 0.05. Micron bar represents 20 μm.

### Inhibition of YAP activity by Verteporfin (VP) affected bovine GCs growth and viability

In order to determine whether VP affects bovine GCs viability, we treated bovine GCs with 5uM VP or vehicle control (M199) for 48 h. FSH showed no survival advantage on bovine GCs growth at 100ng/ml compare to vehicle treated control cells (P > 0.05; Figure 5). VP with FSH treatment still showed 43% inhibition compare to FSH alone. Western blot analysis revealed that TGFa elevated Cyclin D1 protein level (Figure 5A). VP treatment reduced basal as well as TGFs induced Cyclin D1 protein level. FSH has no effect on either basal or VP-reduced Cyclin D1 protein level.

**Figure 5.**
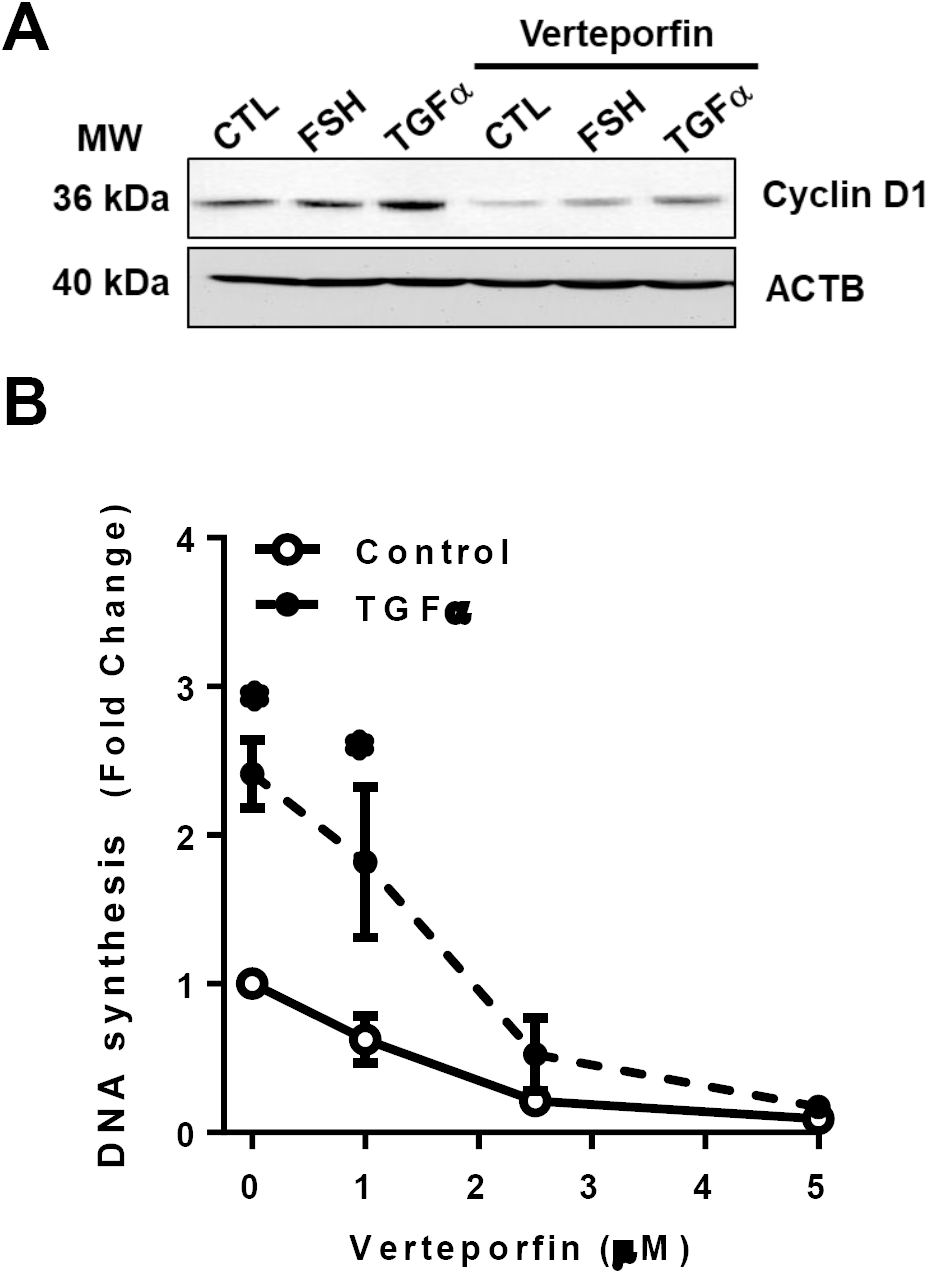
Effects of Verteporfin on cell proliferation in granulosa cells (GC). Verteporfin was used to inhibit Yes-associated protein (YAP1) to determine the effects of YAP on cell proliferation in cultured bovine granulosa cells (GC). Panel A: Representative western blot analysis showing Cyclin D1 expression in cells treated with follicle stimulating hormone (30 ng/mL; FSH) or transforming growth factor alpha (100 ng/mL; TGFα) in the presence of YAP1 inhibitor, verteporfin. Panel B: Quantitative analysis of DNA synthesis in cells treated with increasing concentrations of Verteporfin (0, 1, 2.5. and 5 μM) following treatment of TGFα (100 ng/mL). Data are represented as means ± standard error. ^*^Differ significantly within treatment groups, as compared to control, *P* < 0.05.

### Effects of knockdown of Yes-associated protein (YAP1) on TGFα-induced proliferation in granulosa cells (GC)

In order to test whether YAP plays a role in cell proliferation, we transiently transfected bGCs with YAP siRNA, siTAZ, siYAP and siTAZ or Cy5-labelled Scramble siRNA (siGLO) and treated cells with TNFa for 48 h. Our results showed that siYAP significantly attenuated cell proliferation at basal condition by 46% (P<0.05, Figure 6). TGFa (50 ng/ml) treatment for 48 h significantly increased bGCs proliferation by 2.4 folds (P<0.001, Figure 6). SiYAP however, inhibited TGFa–induced bGCs proliferation by 46% (P<0.001), Figure 6). Western blot analysis indicated that siYAP significantly reduced YAP protein level.

**Figure 6.**
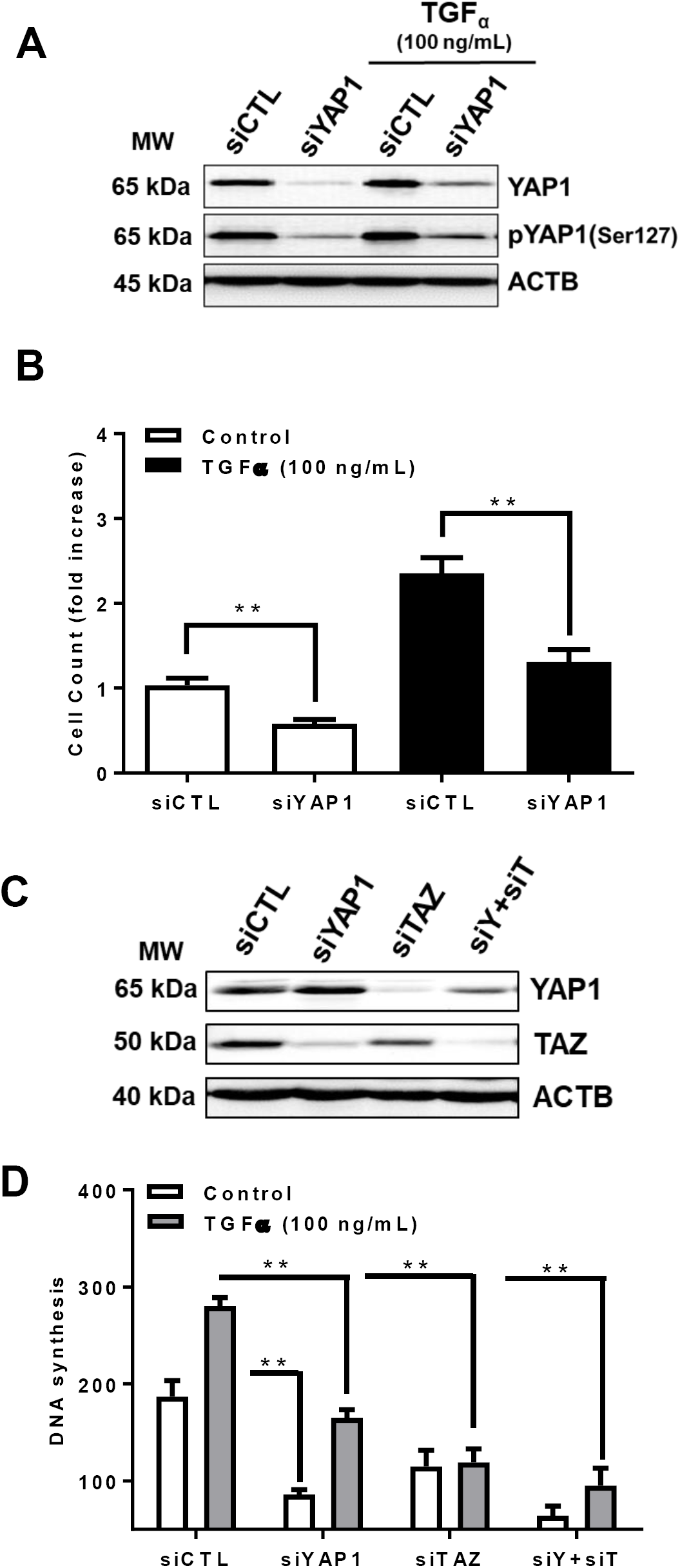
Effects of knockdown of Yes-associated protein (YAP1) on TGFα-induced proliferation in granulosa cells (GC). Yes-associated protein (YAP1) and Tafazzin (TAZ) mRNA were silenced using siYAP1 and siTAZ in bovine granulosa cells (GC). Panel A: Representative western blot showing phospho-YAP1(Ser127) and YAP1 protein expression in siGlo (siCTL) or siYAP1 knockdown GCs, following treatment with transforming growth factor alpha (100 ng/mL; TGFα). Panel B: Mean cell count of siCTL or siYAP1 knockdown cells following treatment with control (n = 3; open bars) or TGFα (n = 3; closed bars). Panel C: Representative western blot showing YAP1 and TAZ total protein expression in siCTL, siYAP1, siTAZ, or a combination of siYAP1 and siTAZ (siY+siT) knockdown GCs. Panel D: Quantitative analysis showing DNA synthesis for siCTL, siYAP1, siTAZ, or siY+siT knockdown cells treated with control (CTL; n = 3; open bars) or TGFα_3_ (n = 3; grey bars). Beta-actin (ACTB; loading control). Data are represented as means ± standard error. ^*^Significant differences between treatment groups, as compared to control, *P* < 0.05.

**Figure 7:**
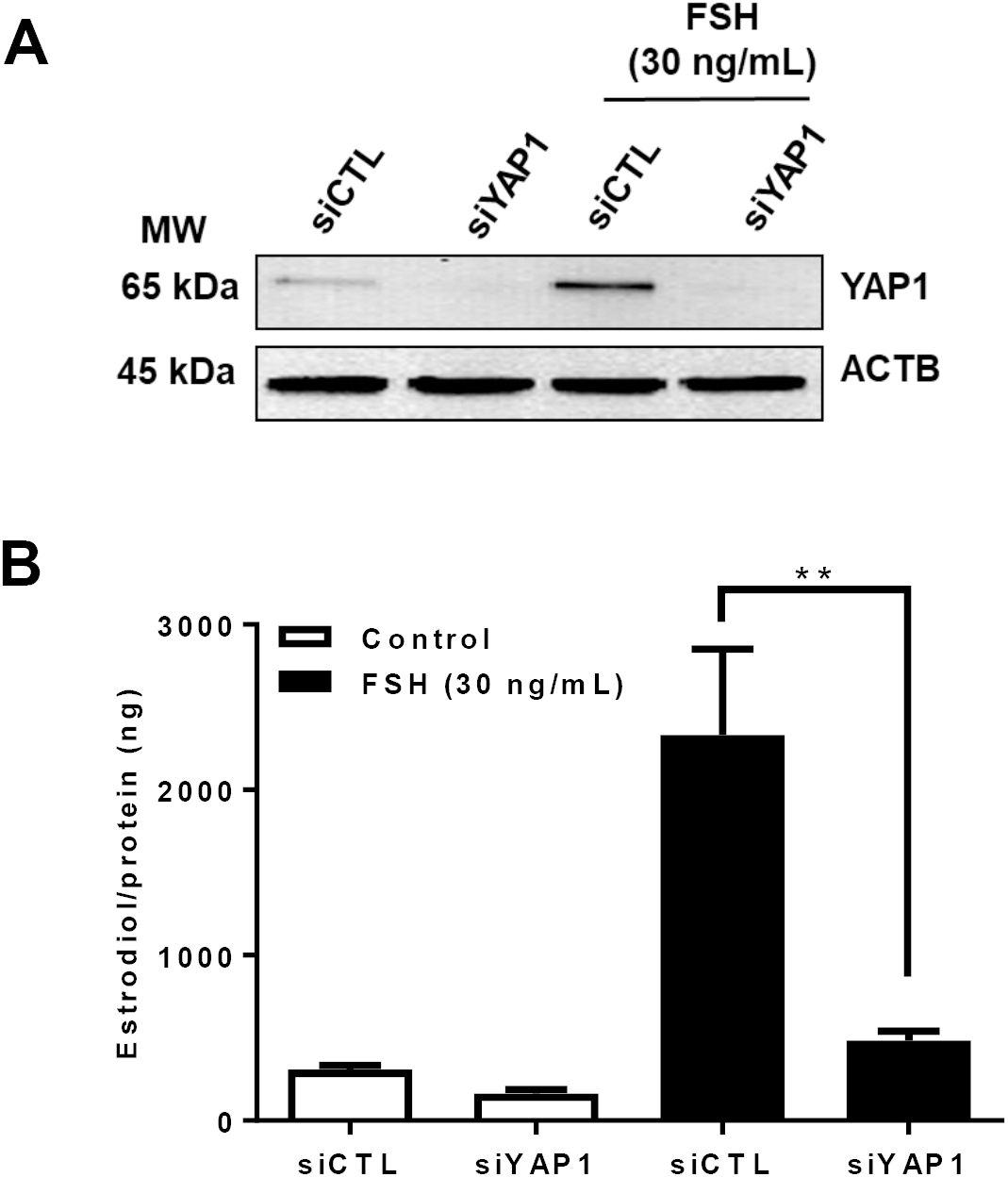
Effects of Yes-associated protein (YAP1) on follicle stimulating hormone (FSH)-induced estradiol production in bovine granulosa cells (GCs). Yes-associated protein (YAP1) mRNA was silenced using siYAP1 to determine the influence of YAP1 on follicle stimulating hormone (FSH)-induced estradiol production in bovine granulosa cells (GCs). Panel A: Representative western blot showing YAP1 total protein expression in siGlo (siCTL) or siYAP1 knockdown GCs, 48-h post-treatment of FSH (30 ng/mL). Panel B: Quantitative ELISA analysis of estrodiol collected from spent media obtained from siCTL (n = 3) or siYAP1 (n = 3) GCs 48-h following treatment with FSH. Beta-actin (ACTB; loading control). Data are represented as means ± standard error. **Significant differences between treatment groups, as compared to control, *P* < 0.05.

### Effects of Yes-associated protein (YAP1) on follicle stimulating hormone (FSH)-induced estradiol production in bovine granulosa cells (GCs)

Granulosa cells uniquely produce estradiol for normal reproductive cycle. FSH is a known hormone to stimulate estradiol production. Here we shown that FSH treatment successfully induced estradiol production by 7.6 folds in bovine GCS (Figure 5, P<0.001). YAP knock down attenuated FSH-induced estradiol production by 80% (Figure 5, P<0.001). Western blot analysis revealed that YAP protein was successfully knockdown.

## DISCUSSION

Hippo signaling is well-established as a regulator of ovarian function and homeostasis, as well as a key mediator and therapeutic target in ovarian cancer cells. However, little is known about the role of YAP in normal ovarian follicle development. Understanding the influence of FSH and ovarian growth factors, such as TGFα, on hippo signaling may lead to better comprehension of the molecular mechanism of proliferation and apoptosis in granulosa cells. Therefore, the present study was undertaken to determine the role of hippo signaling, specifically downstream effector, YAP, on granulosa cells proliferation and steroidogenesis.

Recently, hippo signaling has emerged into a novel molecular target for the regulation of ovarian function and remodeling. Initial work regarding the components of hippo signaling in the ovary were done utilizing Drosophila. Hippo signaling was acknowledged as a key regulator of cell proliferation, migration, and differentiation homeostasis, as well as apoptosis. Recently, studies have emerged establishing the importance of hippo signaling in mammalian tissues, including the ovaries. In the mouse, manipulation of YAP expression regulates ovarian germline stem cells proliferation and differentiation, as well as ovarian function [23]. Disruption of hippo pathway leads to nuclear import and accumulation of YAP and TAZ, co-activating with TEAD to express growth factors [18, 24]. During early embryonic development, deletion of maternal YAP leads to delay in activation of zygotic gene expression; while in wild-type animals, YAP activator, lysophosphatidic acid, can substantially improve early development [25]. In humans, Hippo pathway is a key regulator of granulosa cell proliferation and oocyte maturation. Additionally, as seen in other species, disruption of hippo core elements leads to ovarian genetic abnormalities. In the bovine, nuclear YAP regulates hypoblast embryonic differentiation and inhibition of YAP activity reduces the percent of zygotes that became blastocysts [26]. To our knowledge, this is the first study to examine the role of YAP in bovine granulosa cells.

The components of hippo signaling in granulosa cells have been reported in both the mouse [18], hen [27], and human [28]. Moreover, microarray studies of bovine granulosa cells have revealed the transcripts for hippo signaling [29, 30]. Here, we examined YAP and TAZ protein expression in bovine ovaries. Immunohistochemistry revealed that both YAP and TAZ are expressed in granulosa and theca cells, as well as the corpus luteum. Moreover, we found that YAP is abundant in granulosa cells obtained from follicles ranging from 2-10 mm; however, TAZ only appeared to be highly abundant in granulosa cells obtained from follicles ranging in size from 2-5 mm. Likewise, both YAP and TAZ are expressed in the small and large steroidogenic luteal cells. YAP plays a critical role in the Hippo pathway to regulate cell proliferation in response to cell contact and cell density [31, 32], achieving a density-dependent control of cell proliferation. Granulosa cells were seeded at increasing cell densities of 0.125-0.50 × 10^6^ cells/well to determine how cell contact/density influenced localization of YAP and TAZ. Here we show that when cells seeded at the lowest density had 79.6% greater nuclear YAP then when compared to cells plated at 0.50 x 10^6^ cell/well. Similar to nuclear YAP, cells plated at the lowest density had 67.8% more nuclear TAZ when compared to highest density cells. Our data agrees with other studies demonstrating that cell density influences the localization of YAP and TAZ [33-35].

The D-type Cyclins (Dl, D2 and D3) are a member of the cyclin family which form cyclin-dependent kinase complex with CDKs to governor the cell-cycle Gl phase of the mammalian cell cycle. Cyclin intracellular concentrations are regulated by mitogenic signals such as Ras/MAP kinases, as well as Phosphoinositide 3-kinase. In the bladder [36], Oral squamous [37], gastric [38] cancer cells nuclear YAP leads to the upregulation of Cyclin D1, promoting cell proliferation. In ovarian granulosa cells, cyclin D2 is induced and under the control of FSH signaling, via a cyclic-AMP-dependent pathway [39]. Ovarian thecal cells produce transforming growth factor α (TGFα), which can regulate granulosa cell growth and differentiation in many species including the bovine [40] [41], monkey [42], mouse [43], and human granulosa-like tumor cells [19]. Here we show that inhibition of YAP using Verteporfin decreases the expression of Cyclin D1, regardless of stimulation with FSH or TGFα. Likewise, inhibition of YAP attenuated both basal and TGFα-induced DNA synthesis in granulosa cells.

In mammals, FSH released from the anterior pituitary gland, binds to receptors on granulosa cells, stimulating the expression of CYP19A1, an enzyme responsible for estradiol biosynthesis [1]. Estradiol produced from granulosa cells promotes oogenesis and follicular development. Recently, hippo core components have been reported to influence steroidogenesis in ovarian cells [19]. For instance, an upstream regulator of YAP, SAV1, has been shown to negatively regulate mRNA associated with ovarian follicular steroidogenesis including FSHR and STARD1 [27]. Moreover, knockdown of YAP reduced FSH-induced aromatase (CYP19A1) protein expression in human granulosa-like tumor cells [44]. Additionally, knockdown of YAP reduced FSH-induced estradiol biosynthesis both in vitro [44] and in vivo [23]. Here we show knockdown of YAP inhibited FSH-induced aromatase (CYP19A) expression by 58% when compared to control. Moreover, we demonstrate knockdown of YAP using siRNA attenuates the stimulatory effect of FSH on estradiol biosynthesis in granulosa cells. While hippo signaling is well-established as a regulator of cell proliferation, this data demonstrates hippo signaling, specifically YAP, plays a key role in ovarian steroidogenesis. More studies are warranted to understand the mechanism by while knockdown of YAP attenuates FSH-induced steroidogenesis in granulosa cells.

Following ovulation, luteinizing hormone released from the anterior pituitary gland causes the theca and granulosa cells of the ovulated follicle to differentiate into small and large steroidogenic luteal cells, respectively [45-47]. The luteinized theca and granulosa cells are unique from the undifferentiated theca and granulosa cells in that they are terminally differentiated and do not undergo cell proliferation but rather undergo hypertrophy. We show YAP and TAZ are expressed in luteal tissue, including the small and large luteal cells. However, it is still unknown the role hippo signaling plays in the regulation of luteal cells. In recent studies, overexpression of YAP was shown to induce hypertrophy in skeletal muscle fibers [48, 49], which may also be occurring in luteal cells. Moreover, luteal cells are steroidogenic cells, synthesizing and secreting progesterone, which is require to for the establishment and maintenance of pregnancy. Hippo signaling may be involved in regulating steroidogenesis, as seen in the both current studies and other in ovarian cells [19], however, more work is warranted to determine the role hippo signaling plays in the regulation of luteal cells.

Hippo signaling plays a critical role in maintaining ovarian tissue growth and homeostasis. On the contrary, disruption in hippo signaling leads ovarian abnormalities, tumor progression, and impaired fertility. However, the underlying mechanisms for these disruptive processes remains unclear. Despite remarkable advances in the understanding of hippo signaling and identification of hippo signaling components, regulation of upstream components still requires attention. In the normal functioning ovary, we demonstrate that downstream hippo signaling effector protein, YAP and transcription co-activator, TAZ, are present and localization of both YAP and TAZ are density-dependent. Likewise, YAP and TAZ are critically involved in granulosa cell proliferation. Furthermore, reducing YAP in granulosa cells inhibits FSH-induced aromatase expression and estradiol biosynthesis. The data suggest that YAP plays an important role in the development of ovarian follicles and estradiol synthesis, which are necessary for maintaining normal ovarian function.

## ACKNOWLEDGMENT

Disclosure Statement: The authors have nothing to disclose. The authors thank Janice Taylor and James Talaska at the University of Nebraska Medical Center, Advanced Microscopy Core Facility for their assistance with microscopy. The use of microscope was supported by Center for Cellular Signaling CoBRE-P30GM106397 from the National Health Institutes. This project was supported by Agriculture and Food Research Initiative Competitive Grant no. 2017-67015-26450 from the USDA National Institute of Food and Agriculture; NIH grants R01 HD087402 and R01HD092263, the Department of Veterans Affairs; and The Olson Center for Women’s Health.

---

**Supplementary Figure 1.**
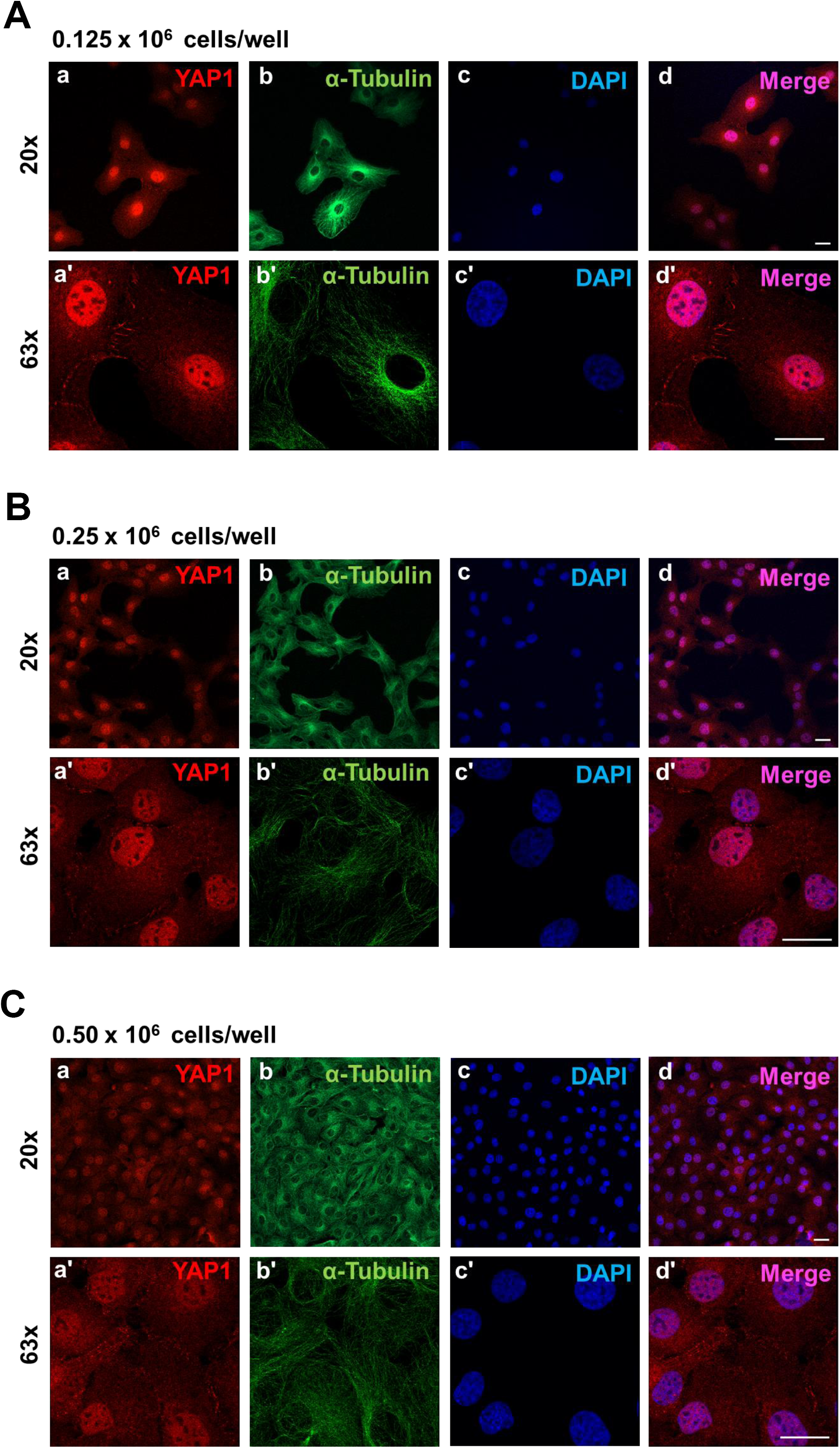
Effects of cell density on localization of Yes-associated protein (YAP1) in bovine granulosa cells (GC). Freshly isolated granulose calls (GC) were seeded in culture dishes at increasing cell densities. Panel A: Representative micrograph of GC plated at 0.125 × 10^6^ cells/well; a) Yes-associated protein (YAP1); 20x magnification, b) alpha-Tubulin; 20x magnification, c) DAPI (nuclei); 20x magnification, d) Merge; 20x magnification, a’) YAP1; 63x magnification, b’) alpha-Tubulin; c’) DAPI; 63x magnification, c’) Merge; 63x magnification. Panel B: Representative micrograph of GC plated at 0.25 × 10^6^ cells/well. Panel C: Representative micrograph of GC plated at 0.50 × 10^6^ cells/well. Micron bar represents 20 μm.

**Supplementary Figure 2.**
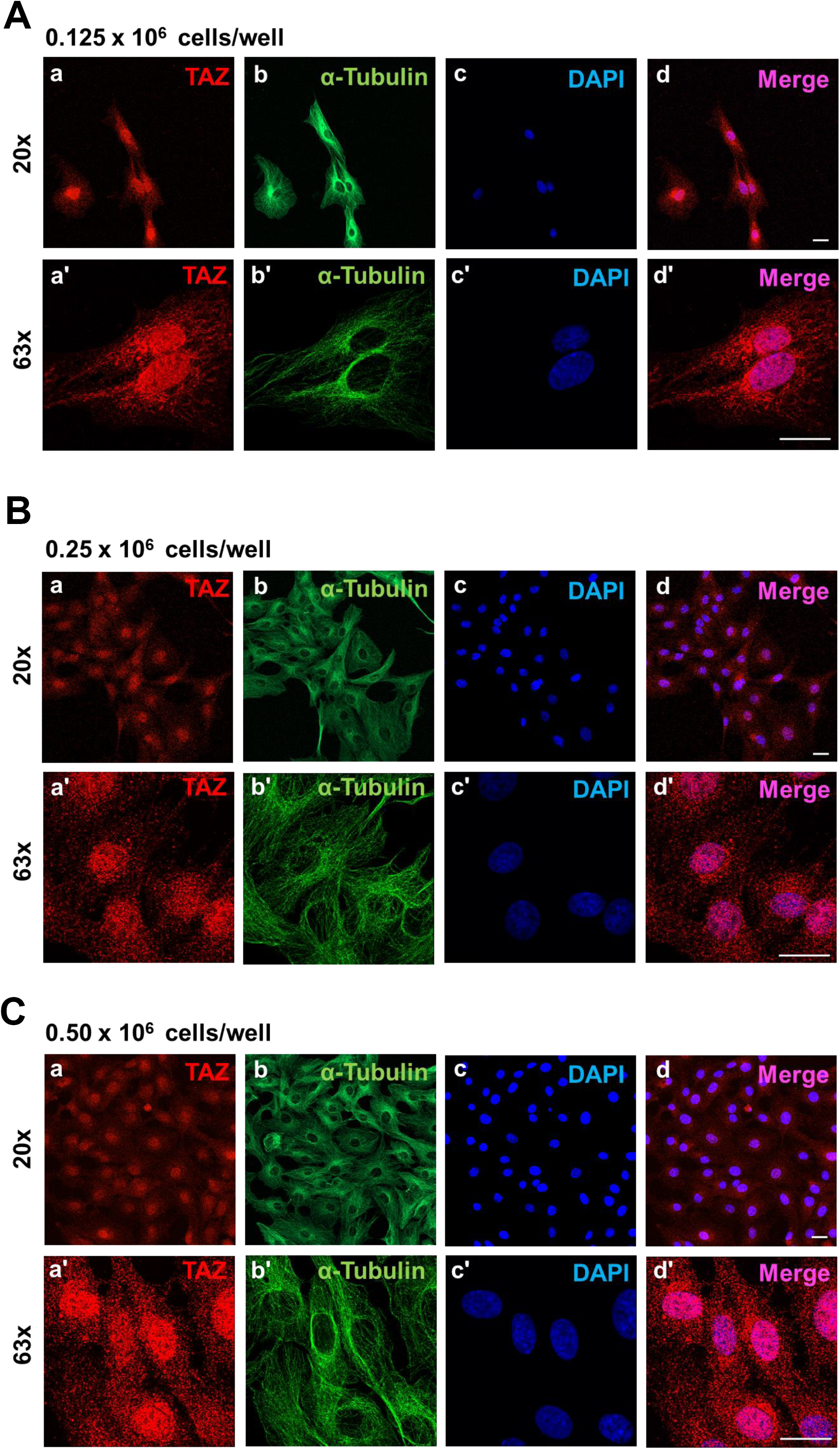
Effects of cell density on localization of Tafazzin (TAZ) expression in bovine granulosa cells (GC). Freshly isolated granulose calls (GC) were seeded in culture dishes at increasing cell densities. Panel A: Representative micrograph of GC plated at 0.125 × 10^6^ cells/well; a) Tafazzin (TAZ); 20x magnification, b) alpha-Tubulin; 20x magnification, c) DAPI (nuclei); 20x magnification, d) Merge; 20x magnification, a’) TAZ; 63x magnification, b’) alpha-Tubulin; c’) DAPI; 63x magnification, c’) Merge; 63x magnification. Panel B: Representative micrograph of GC plated at 0.25 × 10^6^ cells/well. Panel C: Representative micrograph of GC plated at 0.50 × 10^6^ cells/well. Micron bar represents 20 μm.

